# Germline deletion reveals a non-essential role of the atypical MAPK6/ERK3

**DOI:** 10.1101/467597

**Authors:** N. Ronkina, K. Schuster-Gossler, F. Hansmann, H. Kunze-Schumacher, I. Sandrock, T. Yakovleva, J. Lafera, W. Baumgärtner, A. Krueger, I. Prinz, A. Gossler, A. Kotlyarov, M. Gaestel

## Abstract

MAPK6/ERK3 is an atypical member of the MAPKs. An essential role has been suggested by the perinatal lethal phenotype of ERK3 knockout mice carrying a lacZ insertion in exon 2 due to pulmonary disfunction and by defects in function, activation and positive selection of T cells. To study the role of ERK3 *in vivo*, we generated mice carrying a conditional *Erk3* allele with exon3 flanked by LoxP sites. Loss of ERK3 protein was validated after deletion of Erk3 in the female germ line using zona pellucida 3 (Zp3)-*cre* and a clear reduction of the protein kinase MK5 is detected, providing first evidence for the existence of the ERK3/MK5 signaling complex *in vivo.* In contrast to the previously reported Erk3 knockout phenotype, these mice are viable and fertile, do not display pulmonary hypoplasia, acute respiratory failure, abnormal T cell development, reduction of thymocyte numbers or altered T cells selection. Hence, ERK3 is dispensable for pulmonary and T-cell functions. The perinatal lethality, lung and T-cell defects of the previous ERK3 knockout mice are likely due to ERK3-unrelated effects of the inserted lacZ-neomycin-resistance-cassette. The knockout mouse of the closely related atypical MAPK ERK4/MAPK4 is also normal suggesting redundant functions of both protein kinases.

## Introduction

Mitogen-activated protein kinase (MAPK) cascades are conserved eukaryote signaling modules where the downstream effector kinases regulate cell proliferation, differentiation and cell death by phosphorylation of protein substrates. MAPK are also regulators in many physiological processes including development and immune response. Multiple MAPKs were described in mammalian cells, which can be divided in five groups, the ‘classical’ mitogen-responsive MAPKs (ERK1 and ERK2), the stress-activated JNKs (JNK1-3) and p38^MAPK^ (α,β,γ,δ), the big MAPK ERK5, and the atypical MAPKs ERK3, ERK4 and ERK7 (1). MAPKs activity is classically regulated by dual phosphorylation on a TXY motif in the activation loop of the kinase by MAPK-kinases.

ERK3/MAPK6/p97^MAPK^ and the closely-related ERK4/MAPK4/p63^MAPK^ are the only two MAPKs carrying long C-terminal extensions and lacking the dual TXY-phosphorylation motif in the activation loop (2-4). Instead of the canonical dual phosphorylation motif, ERK3 and ERK4 contain only a single phospho-acceptor serine (SEG motif) in the activation loop which can be phosphorylated by p21-activated kinases *in vitro* and in transfected cells (5, 6). Apart from this little is known about the mechanisms of regulation, substrate specificity, and the physiological functions of atypical MAP kinases. Especially, mitogen and stress stimuli resulted in only weak phosphorylation at the SEG motif and the only biological relevant regulator of ERK3 and ERK4 identified so far is the MAP kinase-activated protein kinase MK5 (7-10). MK5 forms a complex with ERK3/4 and is phosphorylated at its activating site T182 within this complex. In turn and still in the complex, ERK3 also auto-phosphorylates at various sites in its C-terminal extension. It is suggested that the ERK3/MK5 complex is involved in the regulation of dendrite morphology and septin function (11). However, it is not clear whether and how the productive complex formation between ERK3/4 and MK5 is regulated by extracellular stimuli or whether it just depends on their expression levels.

Additional functions described for ERK3 include its interaction with the cell cycle regulator Cdc14 (12), its contribution to meiotic spindle stability and metaphase-anaphase transition in mouse oocyte maturation (13) and its interaction with the steroid receptor coactivator 3 (SRC-3), an oncogenic protein overexpressed in multiple human cancers (14). ERK3 seemingly phosphorylates SRC-3 at S857 and regulates it interaction with the ETS- and SP1-type transcription factors (14, 15). Recently, the tyrosyl DNA phosphodiesterase 2 (TDP2), which repairs topoisomerase 2-linked DNA damage, was also described as a substrate of ERK3, and it was suggested that ERK3 phosphorylates TDP2 at S60 and stimulates its phosphodiesterase activity during the DNA damage response (16).

A major contribution to the understanding of the functions of ERK3 and ERK4 was the generation of constitutive knockout alleles of ERK3 (17) and ERK4 (18). Although both kinases are similar in structure and display similar molecular interactions, the phenotypes of both kinase knockouts differed significantly. While ERK4 knockout mice appeared normal, ERK3 knockout mice were not viable, displayed retarded intrauterine growth and pulmonary hypoplasia leading to acute perinatal respiratory failure (17). Furthermore, T cell development, selection and activation was impaired only in the ERK3, but not in the ERK4 knockout mice (19-21). The lack of phenocopy between both knockouts suggested distinct and non-redundant functions of the ERK3/MK5 and ERK4/MK5 signaling complexes.

The perinatal lethality of the constitutive ERK3 knockout mice limits the use of this mouse strain in disease models, which are mostly established for adult mice. To overcome this limitation we generated a conditional allele where exon 3 of *Erk3* is flanked by loxP sites. Here we describe the unexpected finding that germ line deletion of exon 3 in mice causes the complete loss of ERK3 protein but does not lead to any of the phenotypes described for the ERK3 knockout mouse carrying the lacZ insertion. Mice lacking ERK3 protein are viable allowing for the further analysis of ERK3 function in post-natal development and adult mice.

## Materials and Methods

### Generation of ERK3 (MAPK6) conditional knockout mice

*Erk3^ex3lox^*;mice (Mapk6^tm1Mgl^) were generated as indicated in Figure 1A. Briefly, the targeting vector containing lox sites flanked exon3 of mouse *Erk3* gene and FRT sites flanked neomycin cassette was linearized with AsiSI and electroporated in 129Ola ES-cells. Two positive clones (2H3 and 2B5) were obtained by PCR screen and homologous recombination was confirmed by Southern blot analysis. DNA samples were digested with ScaI and probed with a 5’ external probe (PCR product, that amplified 846 bp genomic fragment 5’probe-ERK3-FW: 5’-GTACAGACATGCCTGTACTCATGC-3’ and 5’probe-ERK3-RC: 5’-CTATGCTAACCGACTTAACATGGGAC-3’). Positive clones were injected into blastocysts for the generation of chimeric mice. Agouti germ line pups were derived from the mating of chimeric male mice, obtained following the blastocyst injection of *Erk3* targeted ES cell clone 2H3, with C57Bl/6 Flip females. The resulting *Erk3^ex3loxNeo^*; mice were crossed with C57BL/6-(C3)-Tg(Pgk1-FLPo)10Sykr/J Flippase - expressing mice (22) to delete the neomycin cassette retaining the lox-P-flanked (floxed) exon3 leading *toErk3^ex3lox^*; mice. Subsequent Cre-recombinase expression will then catalyze exon3-excision resulting in an additional frameshift mutation downstream to this exon. For generation of Oocyte specific knockout animals, *Erk3* homozygous floxed mice were crossed with B6-Zp3Cre^tmTgCre^(23). *Erk3^wt/ex3lox^*;:Zp3Cre mice were bred to generate *Erk3^wt/Δex3^*;mice. *Erk3^wt/Δex3^mice* were crossed and littermates of different resulting genotypes (*Erk3*^wt/wt^ (+/+),*Erk3^wt/Δex3^* (+/-) and *Erk3^Δex3/Δex3^* (-/-)) were analyzed. All mice were maintained at the animal facility of the Hannover Medical School under individually ventilated cages (IVC) conditions with free access to food and water. Mice were handled according to the European guideline (2010/63/EU) as well as the German Animal Welfare Act. All animal experiments were approved by the Lower Saxony State Office for Consumer Protection and Food Safety (file Z2017/47).

**Figure 1:**
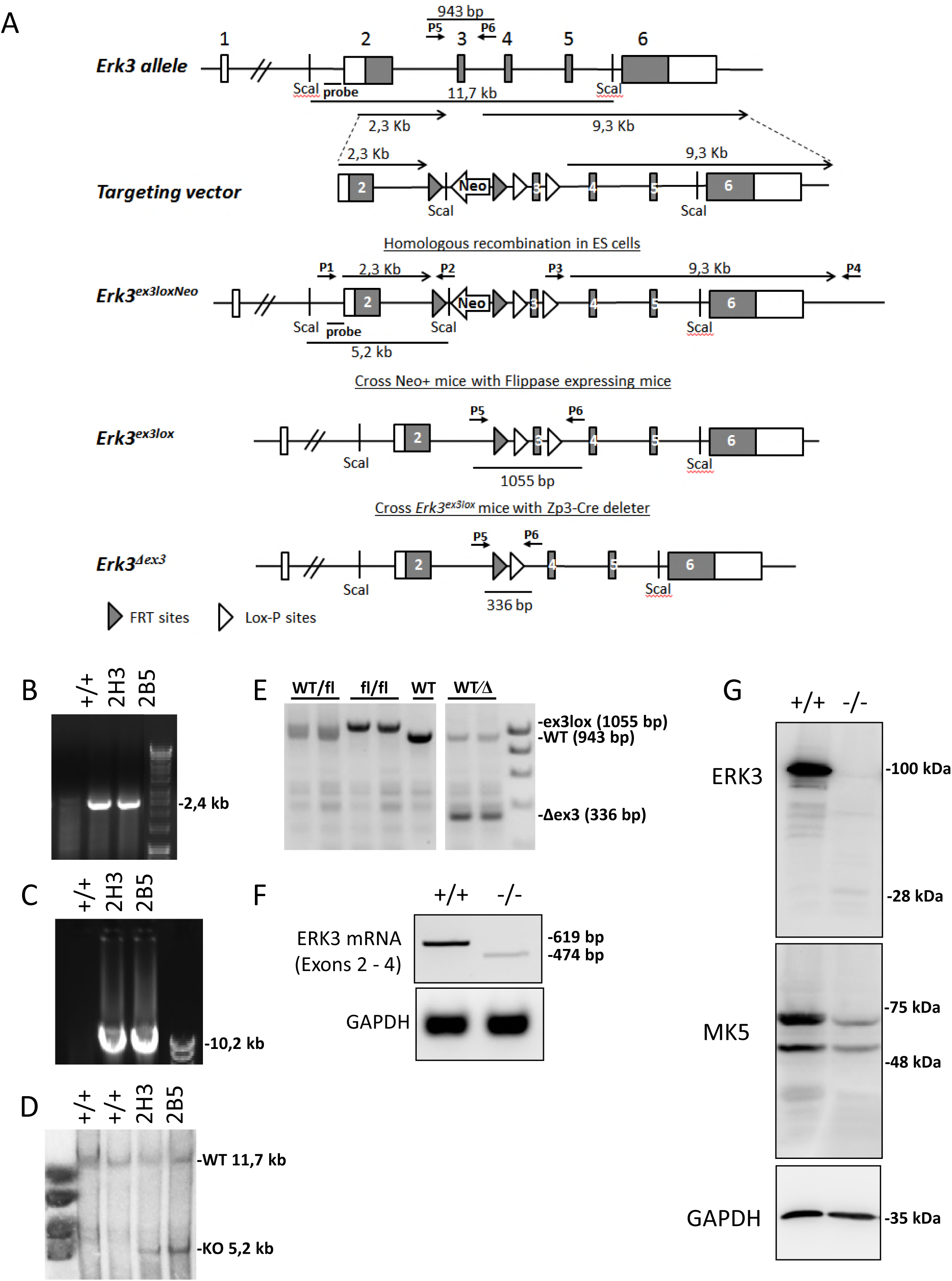
Generation of the conditional ERK3 knockout mouse and deletion of ERK3 mRNA and protein. A) Targeting strategy. B-D) ES cell screening by PCR (B,C) and Southern hybridization (D). B,C) PCR using primer combination P1/P2 and P3/P4 (cf. A) to detect *Erk3^ex3loxneo^* allele. D) Southern blot analysis using probe (cf. A) and ScaI digested DNA. E) Detection of Flp- and Cre-mediated recombination by PCR using primer combinations (P5,P6) leading to fragments as indicated in A. F) ERK3 mRNA was amplified by PCR from total RNA of BMDMs using primers for exons 2 and 4. The ERK3 KO displays a single band weaker and smaller than WT indicating loss of exon 3. G) Western blot analysis of total protein of BMDMs by an N-terminal-ERK3 antibody (Abcam 53277) and a MK5 antibody. Equal loading is demonstrated by GAPDH detection.

### DNA isolation and genotyping

Tail biopsies, cells and colonies were overnight digested at 55°C in lysis buffer (50 mM Tris-Cl (pH 8.0), 100 mM EDTA, 100 mM NaCl and 1% SDS) containing proteinase-K (0.5 mg/mL). For tissue samples proteins were salted out with extra NaCl. DNA was precipitated with isopropanol, washed with 70% ethanol and dissolved in water. Genotyping PCR was performed with Hotstar Taq (Qiagen) with extra Mg2+ under standard conditions. The primers used were: 1) ERK3-1-genotyping-FW: CCGTTTGAGTTTCTTGAGTG, 2) ERK3-3-genotyping-RV: CGTGGTATCGTTATGCG. 1+2 primer combination amplifies 2,4 kb fragment in case of homology recombination (short arm integration).

3) long arm FW: CAGCTTTTGTTCCCTTTAGTGCTCGAC, 4) long arm RC: AGGACTCCTACATCCTGAGCTACCTCTCTAG. 3+4 primer combination amplifies 10,2 kb fragment in case of homology recombination (long arm integration).

5) ERK3_1-target-seq.-FW: TGGACAGAGCACTGGAAG, 6) ERK3-loxP-RC: CTTAAGACAGGAGTGTGGATC. 5+6 primer combination amplifies 943 bp of WT, 1055 bp of exon3 floxed or 336 bp of exon3 deleted fragment.

PCR reactions were separated on 2% agarose gels and images acquired using INTAS Gel documentation system.

### Cell culture

To generate bone marrow-derived macrophages (BMDM), bone marrow cells were flushed from the femurs of mice. Cells were cultured on 10 cm dishes in DMEM supplemented with 10% fetal bovine serum, penicillin/streptomycin, and 50 ng/ml recombinant macrophage colony-stimulating factor (M-CSF) (Wyeth, Boston, MA) under humidified conditions with 5% CO2 at 37°C for 7 days.

### Analysis of ERK3 mRNA expression

Total RNA was isolated from BMDM of ERK3+/+ and ERK3-/- mice. RNA was purified using the Extractme Total RNA extraction kit (BioScience) according to the manufacturer’s instructions. cDNA from 500 ng RNA was synthesized using the first strand cDNA synthesis kit (Fermentas/Thermo) in combination with random hexamer primers. Sense and antisense oligonucleotides (ERK3-mRNA-FW: 5’-TGGACAGAGCACTGGAAG -3’ and ERK3-mRNA-RC: 5’-CTTAAGACAGGAGTGTGGATC-3’) specific for *Erk3* were used to amplify a 619 bp ERK3 mRNA fragment spanning exons 2-4 from ERK3+/+ or a 474 bp fragment lacking the 145 bases of exon 3 from ERK3-/- cells. Amplification of GAPDH mRNA fragment was used as a loading control (GAPDH-fw: 5’-CATGGCCTTCCGTGTTCCTA-3’; GAPDH-rc: 5’-CCTGCTTCACCACCTTCTTGAT-3’).

### SDS-PAGE, Western blot and antibodies

Protein extracts were prepared by direct lysis of the cells in the culture plate with 2x Laemmli’s SDS sample buffer. Protein lysates were separated by sodium dodecyl sulfate polyacrylamide gel electrophoresis (SDS-PAGE) on 7.5%-16% gradient gels and transferred by semi-dry blotting to Hybond ECL nitrocellulose membranes (GE Healthcare). Primary antibodies used were: anti-ERK3 [EP1720Y] ab53277 from Abcam, MK5 [HPA015515] from Atlas antibody, GAPDH from Millipore. Secondary HRP-conjugated antibodies (Santa Cruz Biotechnologies) were used. Antigen-antibody complexes were detected with enhanced chemiluminescence (ECL) detection solution (solution A: 1.2 mM luminol in 0.1 M Tris-HCl (pH 8.6); solution B: 6.7 mM p-coumaric acid in DMSO; 35% H_2_O_2_ solution; ratio 3333: 333: 1) using the Luminescent Image Analyzer LAS-3000 (Fujifilm).

### Histology

Mice were euthanized individually with carbon dioxide in a standard mouse IVC. Lungs were harvested and inflation-fixed with 10% neutral buffered formalin after euthanasia. Lung was cut at different levels, processed through a gradient of alcohols and xylene and embedded in paraffin. For histological examination, 2-3 μm thick sections were cut and stained with hematoxylin and eosin.

### Flow cytometry and cell sorting

Monoclonal antibodies specific for CCR7 (4B12), CD4 (GK1.5), CD5 (53-7.3), CD8α (53-6.7), CD19 (6D5), CD25 (PC61.5), CD44 (IM7), CD62L (MEL-14), CD69 (H1.2F3), Foxp3 (MF23), TCRβ (H57-597), TCRγδ (GL3) were used as AmCyan, Brilliant Violet 510 (BV510), BV421, Pacific Blue (PB), eFluor450, fluorescein isothiocyanate (FITC), Alexa488, Alexa647, phycoerythrin (PE), peridinin chlorophyll protein-Cy5.5 (PerCP-Cy5.5), PE-Cy7, APC, APC-Cy7 and were purchased from eBioscience, BD Biosciences, or Biolegend. Cells were acquired using a BD FACSCanto II and data was analyzed using FlowJo software (Tree Star). Discrimination of dead cells were performed using either the Zombie Aqua Fixable Viability kit or 7-amino-actinomycin D according to the manufacturer’s instructions and doublets were excluded.

### Cell preparations

Thymus and spleen were crushed through a 70 μm cell strainer (Corning) to obtain single-cell suspensions. For spleen, red blood cells (RBCs) were lysed using Qiagen RBC Lysis Solution according to manufacturer’s instructions.

### Statistical analysis

All analysis was performed using GraphPad Prism software (version 7). Data are represented as mean plus or minus SEM. Analysis of significance between 3 groups of mice was performed using one-way analysis of variance followed by Tukey’s test.

### Analysis of T cell proliferation

Splenic and peripheral lymph node single cell suspensions were pooled and stained with cell proliferation dye eFluor670 (eBioscience). Staining was performed at a cell concentration of 1×10^7^ cells/ml in PBS with 1.25 μM eFluor670 for 10 min at 37°C. For in vitro activation 0.15×10^6^ labeled cells in 200 μl medium (RPMI-1640 supplemented with 10% FCS, 1% PenStrep, 1.75 μl β-MercaptoEtOH/500 ml medium, and freshly added rhIL-2 100 U/ml) were seeded into coated round bottom 96-well plates. Plates were coated overnight at 4°C with anti-CD3 (clone 17.A2, final concentration 0.5 μg/ml) and anti-CD28 (clone 37.51, final concentration 1 μg/ml) in a volume of 100 μl per well. After 48h incubation at 37°C 5% CO2 cells were splitted 1:2 into non-coated 96-well plates and supplemented with fresh rhIL-2. 72h and 96h after start of the culture cells were harvested, stained with anti-CD4 (clone RM4-5, Biolegend) and anti-CD8b (clone RMCD8, homemade) antibodies, and proliferation was analyzed by flow cytometry. For live dead discrimination DAPI was used.

## Results

### Conditional targeting of the ERK3 allele

The murine *Erk3* gene is comprised of 6 exons spanning 20 kb of genomic sequence (24). Exons 2 to 6 encode the ORF sequence of ERK3. A targeting vector was designed to flank exon 3 of *Erk3* coding for the activation loop and catalytic kinase subdomains VIII-X (amino acids 186-233) with loxP sites in ES cells (*Erk3^ex3lox^*; Fig. 1 A). The Neo selection cassette flanked by FRT sites was inserted in intron 2. The correct integration of this vector was screened by PCR (Fig. 1B,C) and validated by Southern blot analysis (Fig. 1D) of genomic DNA. Germline transmission was obtained with targeted *Erk3^ex3lox^* ES cell clone 2H3, and the neomycin resistance cassette was removed by breeding with Flp-deleter mice (22). Recombination was confirmed by PCR (Fig. 1E).

### Germline deletion of exon 3 of ERK3 leads to viable mice with the complete loss of ERK3 and reduced MK5 expression

To validate that our targeting strategy results in a null allele we first generated a germ line deletion of exon 3 (*Erk3^Δex3^*) by breeding mice with the floxed exon 3 to Zp3-*Cre* mice that express CRE-recombinase in oocytes (23) which eliminates ERK3 in all cells of the body, i.e. generates a constitutive knockout. Unexpectedly and in contrast to the previously described knockout (17) matings of heterozygous *Erk3^Δex3^* mice gave rise to viable homozygous offspring at the expected Mendelian ratio (n=105 of six crosses and 15 litters (Fig. 2A), 25 WT mice, 45 HETs, 35 ERK3 KO, χ^2^=4.04<5.99 (p<0.05)). PCR analysis of bone marrow-derived macrophages (BMDMs) demonstrated absence of WT mRNA in *Erk3^Δe3x^* homozygotes and the presence of a reduced level of mRNA lacking Exon 3 (Fig. 1F). Western blot analysis of BMDM lysates using various antibodies including the N-terminal specific one, which could detect the C-terminal truncations, revealed the lack of ERK3 in homozygous mutant mice (Fig. 1G). Detection of a very faint band at about 28 kDa could represent some remaining rather instable protein fragment of only part of the catalytic domain (subdomains I-VII) encoded by exon 2 (aa 1-185). Deletion of exon 3 leads to a frame shift and immediate translation stop in potential alternatively spliced transcripts of the targeted allele. Interestingly, the protein level of the ERK3 interaction partner MK5 was also deceased in the ERK3 knockout (Fig. 1G).

**Figure 2:**
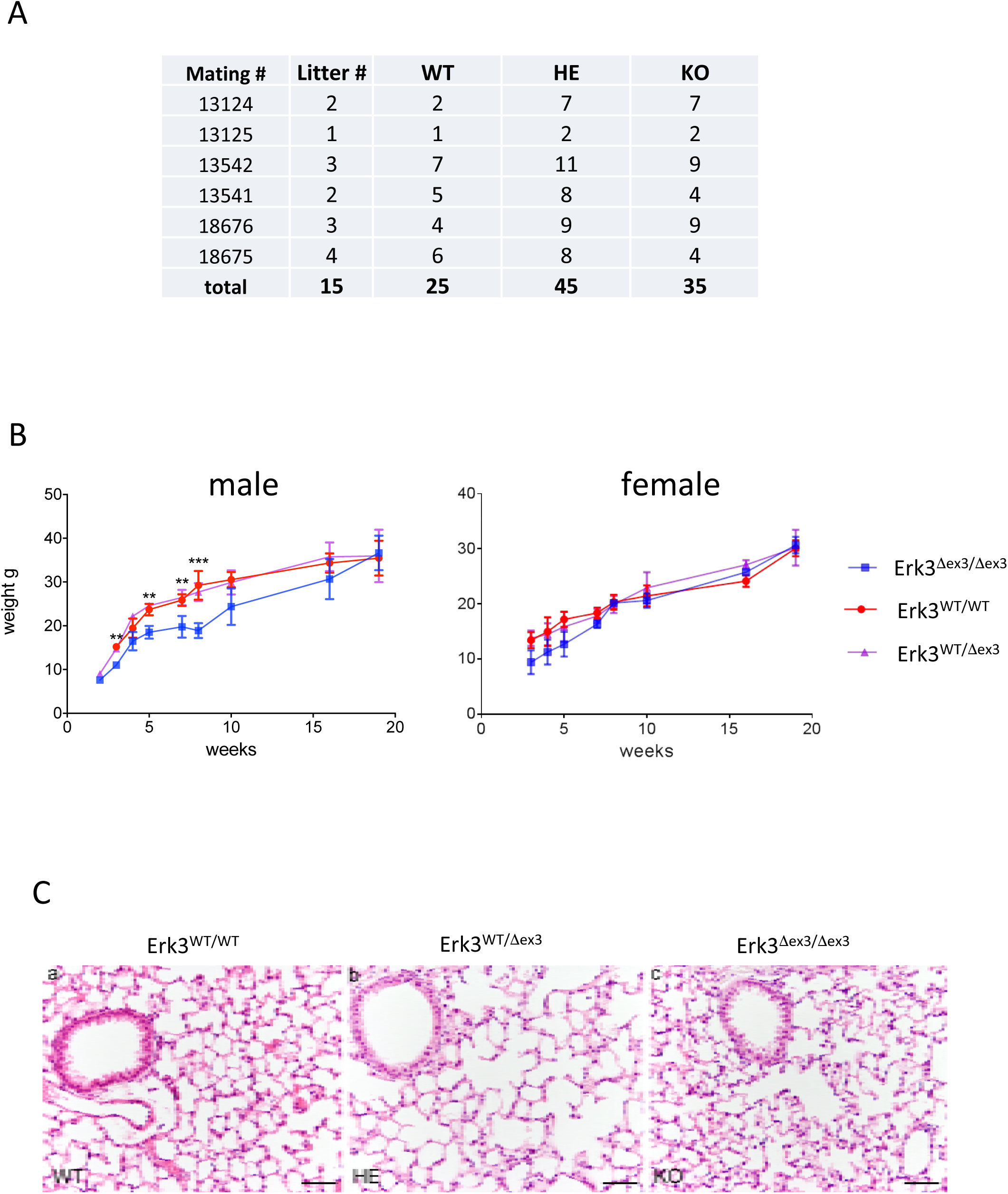
Characterization of the viable ERK3 KO mice. A) Mendelian ratio of the offspring of heterozygous *Erk3^Δex3^* mice. 15 litters of six crosses (total n=105) were analyzed. No statistical significant deviation from Mendelian ratio is detected χ^2^=4.04<5.99 (p<0.05). B) Slight and transient growth retardation of male but not female ERK3 KO mice (4-6 mice per group, ^**^ p<0.05, ^***^ p<0.01). C) Histopathology of the lungs from 35 day old WT (a), ERK3+/- (b) and ERK3 KO (c) mice revealed no significant pathological alterations. Hematoxylin eosin staining, bars = 50μm.

### Early and transient growth retardation of *Erk3^Δex3/Δex3^* mice

We determined body weight of male and female *Erk3^Δex3/Δex3^* mice and their littermates 3 to 20 weeks after birth. Male *Erk3^Δex3/Δex3^* mice displayed significantly lower body weight 3 weeks after birth indicating some early growth retardation (Fig. 2B). However, subsequently all *Erk3^Δex3/Δex3^* mice gradually caught up in weight, developed normally, were apparently healthy and became indistinguishable from wild type and heterozygous littermates in the age of about 20 weeks. Examination of tissue sections stained with hematoxylin and eosin revealed no obvious abnormalities in the lungs of WT, *Erk3^+/Δex3^* and *Erk3^Δex3/Δex3^* mice (Fig. 2C). At present, several mice reached already an age of one year or more, and several males and two females tested were fertile.

### Loss of ERK3 does not impair T-cell development and proliferation

Because of the published effects of the previous ERK3 deletion on T-cell development, activation and function (19-21), we analyzed T-cell development and function in our *Erk3^Δe3x/Δex3^* mice. We first compared T cell populations of the thymus of 5-6 weeks old WT, ERK3+/- and ERK3-deficient mice. We did not detect significant differences in total thymus cellularity. In addition, no differences were observed in the frequencies of any of the major thymocyte populations, CD4 and CD8 double negatives (DN), DP, γδTCR^+^, as well as CD4 and CD8 SP cells (Fig. 3A). This is in contrast to the Erk3-lacZ allele, which strongly reduced numbers of thymocytes, most prominently within the CD4 and CD8 double positive (DP) and CD8 single-positive (SP) subsets (19).

**Figure 3:**
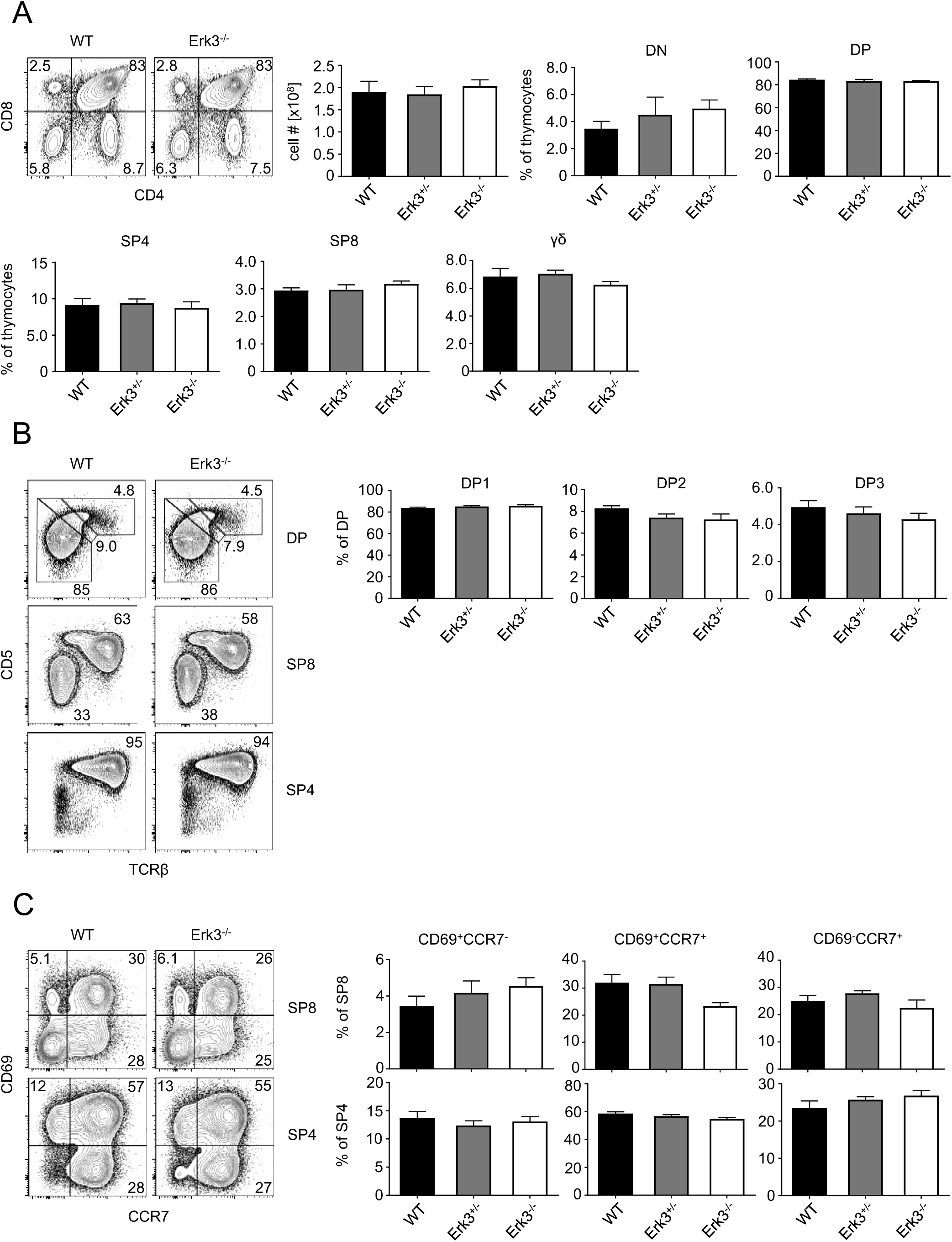
ERK3 is largely dispensable for intrathymic T-cell development. A) Representative flow cytometric analysis of thymi from WT and Erk3^-/-^ mice stained with antibodies against CD4 and CD8α. Numbers in quadrants represent frequencies. Total cellularity of thymi from WT, ERK3^+/-^ and ERK3^-/-^ mice. Statistical analysis of flow cytometric results from WT, ERK3^+/-^ and ERK3^-/-^ mice, cells were defined as DN (CD4^-^, CD8α^-^), DP (CD4^+^, CD8α^+^), SP4 (CD4^+^, CD8α^-^), SP8 (CD4^-^, CD8α^+^) and γδ T cells (CD4^-^, CD8α^-^, TCRγδ^+^). B) Representative flow cytometric analysis of thymi from WT and ERK3^-/-^ mice stained with antibodies against CD4, CD8α, TCRβ and CD5. Numbers adjacent to gates represent frequencies. Statistical analysis of flow cytometric results, cells were defined as DP1 (TCRβ, CD5^lo^), DP2 (TCRβ^int^, CD5^hi^) and DP3 (TCRβ^hi^, CD5^int^) thymocytes. C) Representative flow cytometric analysis of thymi from WT and ERK3^-/-^ mice stained with antibodies against CD4, CD8α, CCR7 and CD69. Numbers in quadrants represent frequencies. Statistical analysis of flow cytometric results, cells were defined as described in A) and CD69^+^CCR7^-^, CD69^+^CCR7^+^ and CD69^-^CCR7^+^ thymocytes. A-C) Pooled data of two independent experiments, n=5-6 per genotype.

It has been proposed that Erk3 contributes to positive selection of thymocytes (21). At steady state, DP thymocytes undergoing selection can be discriminated based on expression of the surface marker CD5 and TCRβ. Pre-selection DP cells are CD5^lo^TCRβ^lo^ (DP1), selecting DP cells are CD5^hi^TCRβ^int^, and CD5^hi^TCRβ^lo^ cells (DP3) are precursors of CD8 SP thymocytes (25). We did not observe any alterations in ratios between the 3 DP thymocyte fractions, suggesting that also positive selection in the thymus is unaffected by loss of ERK3 (Fig. 3B). These data are consistent with normal frequencies of CD4 and CD8 SP cells in the absence of ERK3. In order to assess potential consequences of *Erk3* deletion at the SP stages of T-cell development, we employed the same staining strategy. Again, we found no difference in surface phenotype in CD4 and CD8 SP cells (Fig. 3B). An alternative strategy to monitor thymocyte maturation is staining for the activation marker CD69 and chemokine receptor CCR7. Selection induces upregulation of CD69, followed by CCR7. CD69 is then rapidly downregulated, leaving a mature CD69^-^CCR7^+^ mature thymocyte population. We detected a statistically significant lower frequency of transitory CD69^+^CCR7^+^ thymocytes within the CD8 lineage that had no impact on the frequency of fully mature CD69^-^CCR7^+^ CD8 SP cells in ERK3-deficient thymi (Fig. 3C). We detected no ERK3-dependent differences in frequencies of CD4 SP subsets.

Next, we assessed development of Foxp3^+^ regulatory T cells (Treg) in the absence of functional ERK3. We found no Erk3-dependent differences in Treg cell frequencies in either thymus or spleen (Fig. 4A). The total cellularity of splenocytes was unaltered in the absence of ERK3 (Fig. 4B). Consistent with normal T cell development in the thymus, we also did not detect any alteration in the ratio of T vs. B cells in spleen, although we noted a marginal, but statistically significant increase in the frequency of splenic B cells. In addition, we found no changes in the ratio between CD4^+^ and CD8^+^ T cells upon deletion of ERK3. Expression of CD44 and CD62L allows for discrimination of naïve (CD44^-^CD62L^+^), central memory (CD44^+^CD62L^+^) and effector memory (CD44^+^CD62L^-^) T cell subsets. We found no differences in the frequencies of naive, central memory or effector T cells within the CD4 and CD8 T-cell compartments in spleen when comparing ERK3-sufficient and ERK3-deficient mice (Fig. 4C). These data suggest that loss of Erk3 expression does not result in aberrant T-cell activation at steady-state. T cells from conventional ERK3 KO mice had reduced proliferation capacity upon unspecific TCR stimulation using anti-CD3 and anti-CD28 antibodies (4). Although T-cell activation was not altered in our newly generated ERK3 knockout mice at steady state, we compared proliferation capacity of WT and ERK3-deficient cells upon CD3/CD28 stimulation. In contrast to the conventional ERK3 KO we could not observe any differences in T cell proliferation, neither in CD4 nor CD8 T cells, for our new ERK3 KO (Fig. 4D). Together, we conclude from these data that ERK3 is dispensable for intrathymic T cell development, T cell homeostasis as well as T cell proliferation.

**Figure 4:**
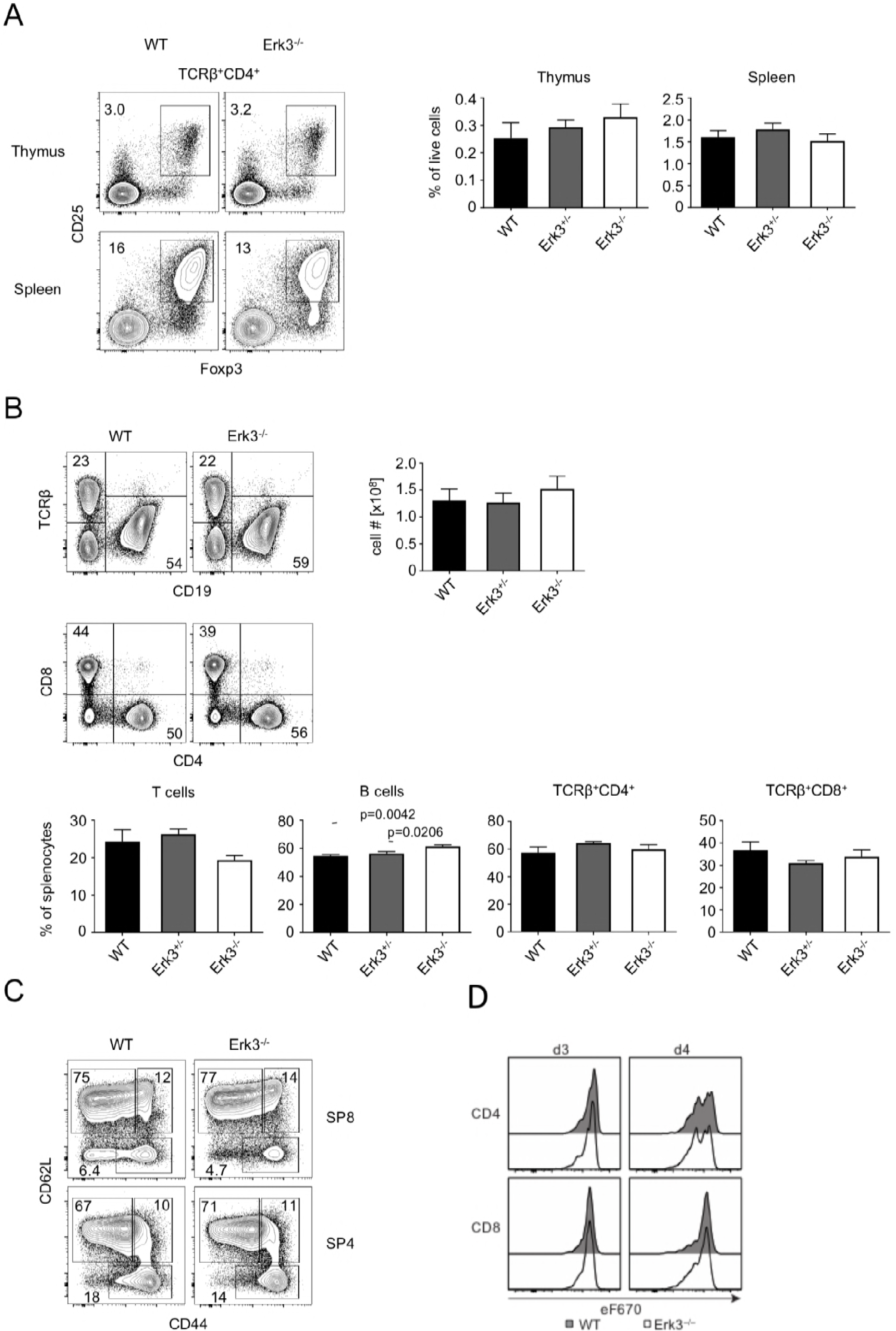
ERK3 is largely dispensable for T-cell homeostasis and proliferation. A) Treg cell numbers in thymus and spleen. Dot plots show representative flow cytometric analysis of thymi and spleen from WT and ERK3^-/-^ mice stained with antibodies against CD4, TCRβ, Foxp3 and CD25. Numbers adjacent to gates represent frequencies. Graphs display statistical analysis of flow cytometric results from WT, ERK3^+/-^ and ERK3^-/-^ mice. B) Representative flow cytometric analysis of spleen from WT and ERK3^-/-^ mice stained with antibodies against CD19, TCRβ, CD4 and CD8. Numbers in quadrants represent frequencies. Total cellularity of spleen from WT, ERK3^+/-^ and ERK3^-/-^ mice. Statistical analysis of flow cytometric results from WT, ERK3^+/-^ and ERK3^-/-^ mice, cells were defined as T cells (TCRβ^+^), B cells (CD19^+^), SP4 (TCRβ^+^, CD4^+^) and SP8 (TCRβ^+^, CD8^+^). C) Representative flow cytometric analysis of SP4 and SP8 splenic T cells from WT and ERK3^-/-^ mice stained with antibodies against CD4, CD8, CD44 and CD62L to identify naïve (CD44^-^CD62L^+^), central memory (CD44^+^CD62L^+^) and effector memory (CD44^+^CD62L^-^) T cell subsets. A-C) Pooled data of two independent experiments, n=5-6 per genotype. D) Flow cytometric analysis of CD4 (upper panel) and CD8 (lower panel) T cell proliferation from WT and ERK3^-/-^ mice after 3 and 4 days of culture. Shown are representative contour plots of 1 – 2 experiments with each n = 2 mice per genotype.

## Discussion

Here, we report the generation and analysis of a novel *Erk3* null allele, and show that deletion of ERK3 is not essential for viability, pulmonary function or T-cell development in contrast to previously described mice carrying a lacZ insertion into exon2 of *Erk3* (17, 19-21). The reason for these differences likely lie in the different targeting strategies used to disrupt *Erk3.* The constitutive ERK3 knockout was generated by insertion of a lacZ in frame with the translation start of exon 2 of the ERK3 gene 12 amino acids downstream of the initiation codon and a neomycin-resistance-cassette downstream of lacZ. The conditional allele reported here removes exon 3 of *Erk3* and leaves only a single FRT and a single loxP sites in intron 2. The insertion of a lacZ-neomycin-resistance-cassette often altered transcription of genes in the flanking DNA around the targeted gene, and accordingly the phenotype of the mutation (26-29). This may be due to the loss or disruption of intragenic regulatory elements, the constitutive promoter driving the neomycin gene, removal of insulating DNA in the targeted alleles, and local silencing due to disruption of normal chromatin organization by the exogenous construct (26). The detailed transcriptome analysis of 29 targeted alleles in mice revealed that down-regulated genes flanking the targets were rather equally distributed 5’ and 3’ of the target and the median distance from the target was 34kb (26). In this regard, it is interesting that on chromosome 9 the gene for an essential subunit of RNA polymerase II, leo1, is 40 kb 3’ to the *Erk3* gene. Hence, one may speculate that down regulation of essential genes, such as leo1, may severely compromise cell viability and contribute to the previously reported *Erk3* mutant phenotype. In contrast to effects of insertion cassettes and to the best of our knowledge no effects of insertions of single loxP and FRT sites into intronic sequences are described. Therefore, a likely explanation for the perinatal lethality, lung and T-cell defects observed in the previous ERK3 knockout mice is ERK3-unrelated effects of the inserted lacZ-neomycin-resistance-cassette.

We and others have described the ERK3/MK5 and ERK4/MK5 signaling modules (7, 9, 10, 30). Deletion of MK5 and ERK4 in mice resulted in some mild growth retardation (31, 32) and no significant phenotype at all (18), respectively. In contrast, disruption of *Erk3* had a lethal phenotype (17). So far, it was difficult to understand that deletion of different components of these modules do not mutually phenocopy, but display completely different phenotypes. This also questioned the physiological relevance of these signaling modules. However, the phenotype of slight and transient male growth retardation described for deletion of exon 3 of *Erk3* here phenocopies the MK5 knockout and reconciles the genetic analysis of these signaling complexes with their existence in a specific signaling module. Furthermore, the observation that the level of the interaction partner MK5 is significantly reduced in the *Erk3^Δex3/Δex3^* BMDMs strongly supports the existence of the ERK3/MK5 signaling complex *in vivo.* Possibly the remaining MK5 in our *Erk3^Δex3/Δex3^* mice is stabilized by its interaction with ERK4.

So far, the differences between the phenotypes of the conventional knockouts of ERK3 and ERK4 where explained by specific non-redundant functions of these closely related atypical MAPKs (18), which contrasts with the similarities in structure and molecular interactions of both kinases (5, 33, 34). The viability and fertility of mice with deletion of exon 3 of *Erk3* is much more similar to the phenotype of the conventional ERK4 knockout. Hence, one cannot exclude a redundant function of both atypical MAPKs, which could be revealed in the future by generation of the exon 3 ERK3/ERK4 double knockout mouse.

## Acknowledgements

The work of M.G. is supported by Deutsche Forschungsgemeinschaft.

